# Predicting functional long non-coding RNAs validated by low throughput experiments

**DOI:** 10.1101/634345

**Authors:** Bailing Zhou, Yuedong Yang, Jian Zhan, Xianghua Dou, Jihua Wang, Yaoqi Zhou

## Abstract

High-throughput techniques have uncovered hundreds and thousands of long non-coding RNAs (lncRNAs). Among them, only a small fraction has experimentally validated functions (EVlncRNAs) by low-throughput methods. What fraction of lncRNAs from high-throughput experiments (HTlncRNAs) is truly functional is an active subject of debate. Here, we developed the first method to distinguish EVlncRNAs from HTlncRNAs and mRNAs by using Support Vector Machines and found that EVlncRNAs can be well separated from HTlncRNAs and mRNAs with 0.6 for Matthews correlation coefficient, 64% for sensitivity, and 81% for precision for the independent human test set. The most discriminative features are related to sequence conservations at RNA (for separating from HTlncRNAs) and protein (for separating from mRNA) levels. The method is found to be robust as the human-RNA-trained model is applicable to independent mouse RNAs with similar accuracy and to a lesser extent to plant RNAs. The method can recover newly discovered EVlncRNAs with high sensitivity. Its application to randomly selected 2000 human HTlncRNAs indicates that a large number of functional lncRNAs are waiting to be validated. The method is expected to speed up and reduce the cost of the discovery by prioritizing potentially functional lncRNAs prior to experimental validation. EVlncRNA-pred is available as a web server at http://biophy.dzu.edu.cn/lncrnapred/index.html. All datasets used in this study can be obtained from the same website.

## Introduction

Advances in high-throughput sequencing and microarray technologies showed that most of the human genome transcribe into RNAs despite they were not coded for proteins [1-3]. Among these non-coding RNAs (ncRNAs), long transcripts (>200 nucleotides) with unknown functions were found prevalent with low expression, highly tissue-specific, and lack of strong cross-species conservation [4-6]. Some long ncRNAs (lncRNAs) have been confirmed as functional and disease-relevant using traditional low throughput techniques such as qRT-PCR, knockdown, Western blot, Northern blot, and luciferase reporter assays [7-11]. So far, more than 1000 lncRNAs in >70 species were experimentally validated and collected in a number of databases (lncRNADisease, lncRInter, lncRNAdb, PLNlncRBase) [12-15]. These databases were integrated into the comprehensive EVlncRNAs database [16], which collected all known EVlncRNAs up to May, 2016 from 77 species.

The experimentally validated lncRNAs (EVlncRNAs), however, are only a tiny fraction of all transcribed ones. What percentage of transcribed lncRNAs is functional remains a subject of active debate [17]. It is known that some lncRNAs can be expressed due to lack of fidelity in transcription initiation by RNA polymerase II [18]. Hon *et al*.[19] found that 69% of 27,919 FANTOM CAT lncRNAs overlap with trait-associated single nucleotide polymorphisms. However, the overlap could be due to their genomic positions, rather than intrinsic functions coded in sequences [20]. Nevertheless, 31% lncRNAs remain unaccounted for. Liu *et al.*[21], on the other hand, found that only 3% (499/16, 401) lncRNA loci are essential for robust cell growth, based on a large-scale knockdown using a CRISPR interference technique. More importantly, analysis of mutational loads suggests that “the functional fraction within the human genome cannot exceed 25% and is probably considerably lower” [22]. Thus, a significant portion of transcribed lncRNAs is possibly non-functional.

Existence of non-functional but transcribed lncRNAs calls for computational methods to prioritize potentially functional lncRNAs prior to expensive and laborious experimental validations. Current computational tools on identification of lncRNAs have been focused on distinguishing expressed lncRNAs from coding RNAs [23], a challenging problem as some lncRNAs were coded for short peptides while others such as H19, Xist, Mirg, and Gtl2 have predicted coding regions of longer than 100 amino acids [24, 25]. One approach for lncRNA identification is cross-species comparison. Examples are CRITICA [26], PhyloCSF [27], CSTMiner [28] and RNAcode [29]. Because the majority of lncRNAs are not conserved across different species, many methods estimated coding potentials of a sequence by using a wide variety of features and machine-learning techniques. Examples are CPAT by logistic regression model [30], lncRNA-ID [31], lncRNApred [32], FEELnc [33] and COME [34] based on random forest and PORTRAIT [35], CNCI [36], CPC [37], PLEK [38] and lncRScan-SVM [39] based on support vector machines. Using high-throughput experimental data have proven useful for further improving the accuracy of discrimination of lncRNAs from mRNAs [34, 40-42].

In this study, we address the question whether or not lncRNAs experimentally validated (EVlncRNAs) by low throughput techniques are distinguishable from those lncRNAs obtained from high throughput experiments (HTlncRNAs). By using support vector machines and employing sequence-derived and HT experimental features in combination or separately, we showed that experimentally validated lncRNAs are identifiable from HTlncRNAs and mRNAs with reasonable accuracy. Moreover, a method trained and tested from human datasets is applicable to mouse RNAs with similar accuracy and to plant RNAs with somewhat lower accuracy. This indicates the robustness of the method developed for locating functional lncRNAs. The online server of EVlncRNA-pred is freely available at http://biophy.dzu.edu.cn/lncrnapred/index.html.

## Results

### Model performance for the full-feature model

Using positive samples collected in the EVlncRNAs dataset [16] and negative samples from lncRNAs and mRNAs from GENCODE [43] (see Methods), we built the training set from human RNAs and independent test sets from human, mouse and plant RNAs. Table 1 shows the results of the human 10-fold cross-validation and independent test by the support vector machines (SVM) model with 33 features (the full-feature model). The corresponding ROC curves are shown in Figure 1. The results indicate that the model performs better on the test set (MCC at 0.60 compared to 0.51, AUC at 0.88 compared to 0.84). This is likely due to a slightly larger training set (799 positive samples) than in the cross validation (719 positive samples). When the model is further applied to mouse RNAs, there is a performance drop to the performance level similar to ten-fold cross validation. It should be noted that the lower precision for the mouse test set is due to higher sensitivity as the threshold was set by the 10-fold cross validation. If one adjusts the threshold to sensitivity of 0.64, one would obtain a precision of 0.74. Nevertheless, the low standard deviation in ten-fold cross-validation performance over 100 randomly selected ten folds and the consistently high, cross-species performance (AUC>0.84) in independent tests confirm the overall quality and robustness of the model developed.

**Table 1.** Performance of full-feature and sequence-only SVM models trained on human datasets

|  |  | MCC | AUC | Accuracy | Sensitivity | Specificity | Precision |
| --- | --- | --- | --- | --- | --- | --- | --- |
| Full-feature model |  |  |  |  |  |  |  |
| Human | CV<sup>a</sup> | 0.513±0.006 | 0.841±0.006 | 0.791±0.003 | 0.599±0.011 | 0.887±0.005 | 0.728±0.005 |
|  | Test<sup>b</sup> | 0.603 | 0.879 | 0.829 | 0.641 | 0.923 | 0.806 |
| Mouse | Test<sup>c</sup> | 0.512 | 0.859 | 0.765 | 0.777 | 0.759 | 0.617 |
| Sequence-only model |  |  |  |  |  |  |  |
| Human | CV<sup>a</sup> | 0.471 | 0.841 | 0.774 | 0.551 | 0.887 | 0.709 |
|  | Test<sup>b</sup> | 0.514 | 0.852 | 0.792 | 0.590 | 0.893 | 0.734 |
| Mouse | Test<sup>c</sup> | 0.481 | 0.846 | 0.745 | 0.777 | 0.729 | 0.589 |

<sup>a</sup> 10-fold cross-validation on the full training set. The mean and standard deviation are obtained from 100 random divisions of 10 folds in the training set. <sup>b</sup> Test results on the independent test set of the human RNAs. <sup>c</sup> Test results on the independent test set of the mouse RNAs.

**Figure 1.**
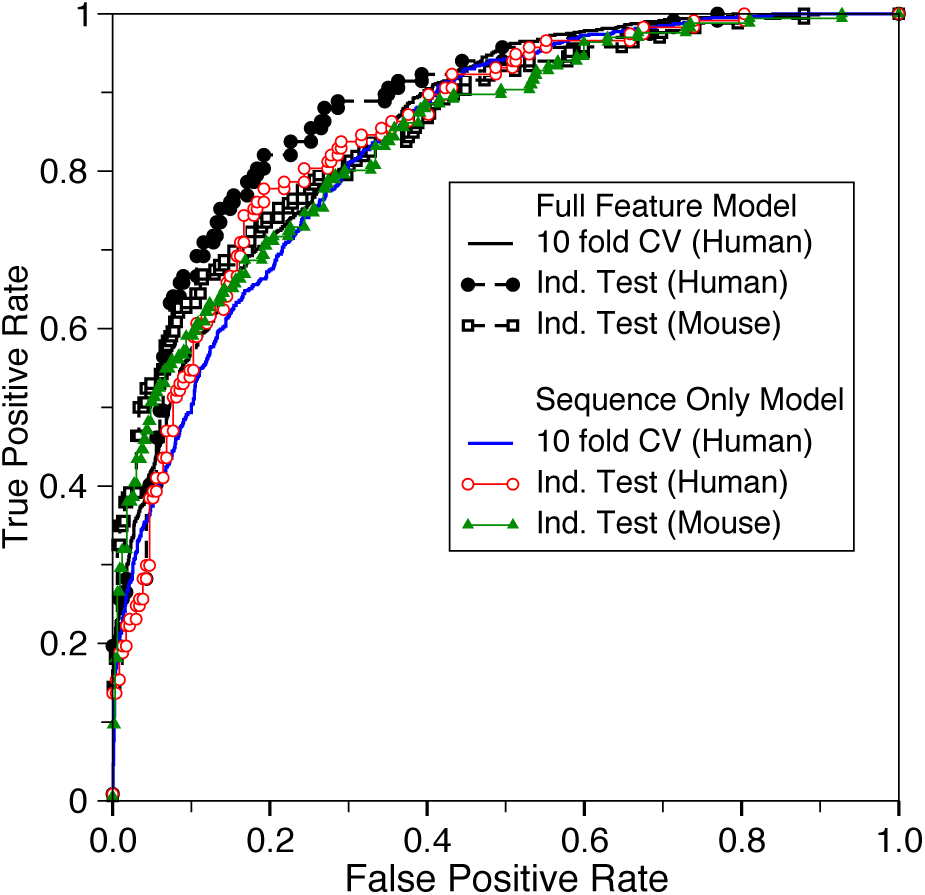
Receiver operating characteristic curves by full-feature and sequence-only models trained on human RNAs.

### Model performance for the sequence-only model

The above model employed some high-throughput experimental results including expression abundance and histone modification. However, these experimental data are not always available. Thus, we also built a model that requires the input of a sequence only. Table 1 and Figure 1 also present the results from the sequence-only model. The model performance is much more similar among the ten-fold cross validation and two independent tests (human and mouse test sets) with MCC=0.47, 0.51, 0.48 and AUC=0.84, 0.85, and 0.85, respectively. This overall performance is slightly worse than the case when experimental data were employed, confirming the usefulness of expression abundance and histone modification in EVlncRNA discrimination. On the other hand, if these experimental results are not available, the sequence-only model yields adequate accuracy in separating EVlncRNAs from HTlncRNAs and mRNAs as shown in Figure 1 and Table 1.

### Model performance for the plant test set

The ability of human-RNA trained model to predict mouse lncRNA indicates inherently similar characteristics of functional lncRNAs in human and mouse. It is of interest to know if plant EVlncRNAs can also be detected in a similar accuracy. In human and mouse, we used phastCons [44] scores provided by the UCSC [45] to represent the DNA sequence conservation. However, UCSC do not have phastCons scores of *Arabidopsis thaliana*. Thus, we re-trained all models without using DNA conservation scores.

Table 2 and Figure 2 compare the performance of the new full-feature and sequence-only models without DNA conservation in 10-fold cross-validation and independent tests on human, mouse, and plant RNAs by models trained on human RNAs. The results show that there is a large drop in performance when the human-RNA-trained model is applied to plant with MCC decreasing from 0.56 to 0.38 and AUC decreasing from 0.87 to 0.80 for the full feature model without DNA conservation. By comparison, the corresponding changes from human to mouse are 0.56 to 0.52 for MCC and 0.87 to 0.84 for AUC. This suggests that the difference between human and plant lncRNAs is larger than the difference between human and mouse lncRNAs. Nevertheless, applying human-RNA trained model to plant remains highly discriminative with AUC=0.80 for the full-feature model and 0.73 for the sequence-only model (all without DNA conservations). This confirms the robustness of the model trained by human RNAs and the existence of basic common characteristics of EVlncRNAs from plant to human. This also suggests that plant-specific training when sufficient data is available may be necessary to maximize the discrimination capability.

**Table 2.** Performance of full-feature and sequence-only SVM models (except DNA conservation scores) trained on human datasets

|  | MCC | AUC | Accuracy | Sensitivity | Specificity | Precision |
| --- | --- | --- | --- | --- | --- | --- |
| Full-feature model except DNA conservation |  |  |  |  |  |  |
| Human-CV <sup>a</sup> | 0.482 | 0.845 | 0.781 | 0.514 | 0.915 | 0.753 |
| Human-Test <sup>b</sup> | 0.560 | 0.869 | 0.812 | 0.581 | 0.927 | 0.800 |
| Mouse-Test <sup>c</sup> | 0.518 | 0.844 | 0.779 | 0.723 | 0.807 | 0.652 |
| Plant-Test <sup>d</sup> | 0.383 | 0.801 | 0.744 | 0.417 | 0.908 | 0.694 |
| Sequence-only model except DNA conservation |  |  |  |  |  |  |
| Human-CV <sup>a</sup> | 0.448 | 0.833 | 0.763 | 0.562 | 0.864 | 0.675 |
| Human-Test <sup>b</sup> | 0.521 | 0.849 | 0.792 | 0.632 | 0.872 | 0.712 |
| Mouse-Test <sup>c</sup> | 0.414 | 0.818 | 0.719 | 0.711 | 0.723 | 0.562 |
| Plant-Test <sup>d</sup> | 0.221 | 0.725 | 0.689 | 0.283 | 0.892 | 0.567 |

<sup>a</sup> 10-fold cross-validation on the full training set. <sup>b</sup> Test results on the independent test set of the human RNAs. <sup>c</sup> Test results on the independent test set of the mouse RNAs. <sup>d</sup> Test results on the independent test set of the plant RNAs.

**Figure 2.**
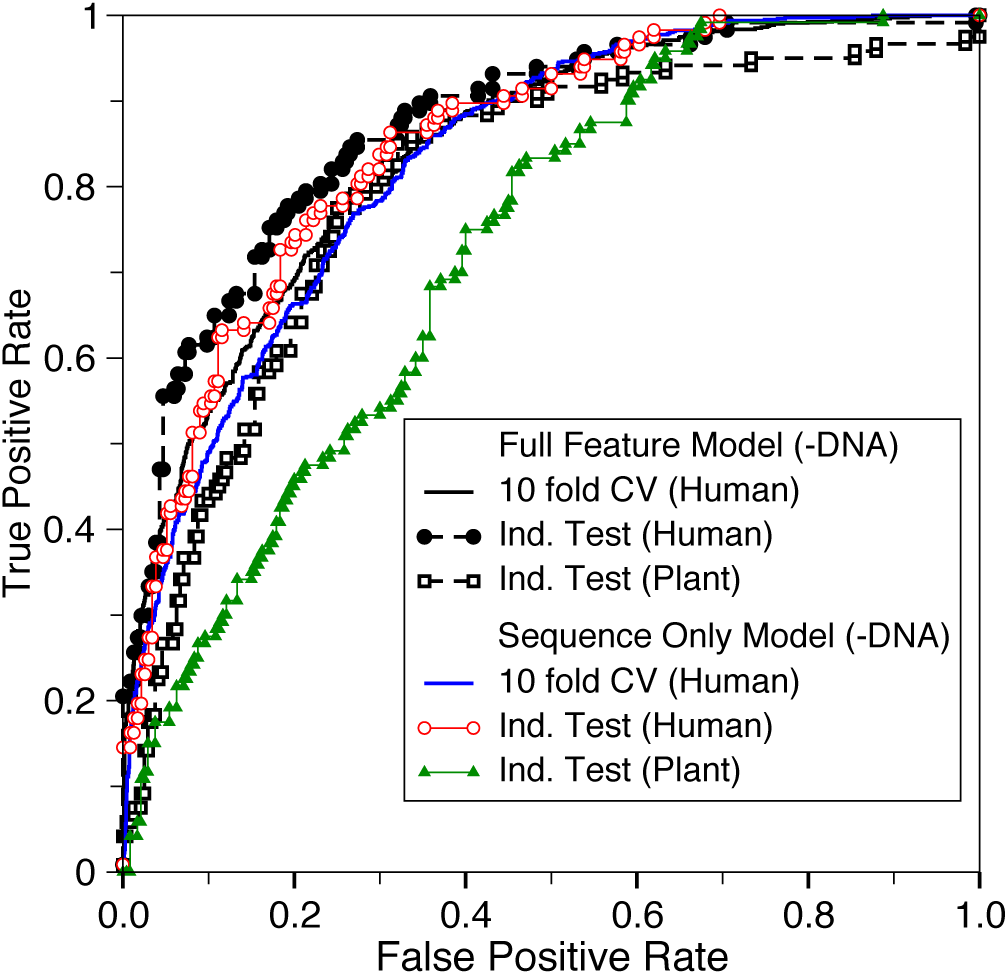
As in Figure 1 but for the model without DNA conservation features and tested by the plant RNAs.

### The importance of individual features

To examine the discrimination power of each feature group for separating EVlncRNAs from HTlncRNAs and mRNAs, we obtained the best performing single feature in each feature group according to 10-fold cross-validation and compare them in Figure 3. For single-feature performance, the performance is measured by the difference between the AUC (ΔAUC) by the single feature group and 0.5 by random prediction or between the AUC by the full feature model and by the model after removing the single feature group. Figure 3 shows that according to feature removal, protein conservation has the highest ΔAUC value from the full feature mode at 0.031, followed by RNA conservation (ΔAUC = 0.020). The best experimental feature is the H3K4me3 modification group with the ΔAUC value at 0.008. These changes in ΔAUC are small. We can also measure the changes in precision at a fixed value of sensitivity. The same trend is observed with two highest reductions of 10.3% and 8% in precision for removing protein and RNA conservation, respectively, and the largest reduction of 3.7% for removing the H3K4me3 modification group in experimental features. This result is similar to analysis of models based on single feature only (Figure 3). All features contribute somewhat to lncRNA classifications except small RNA-seq experimental data.

**Figure 3.**
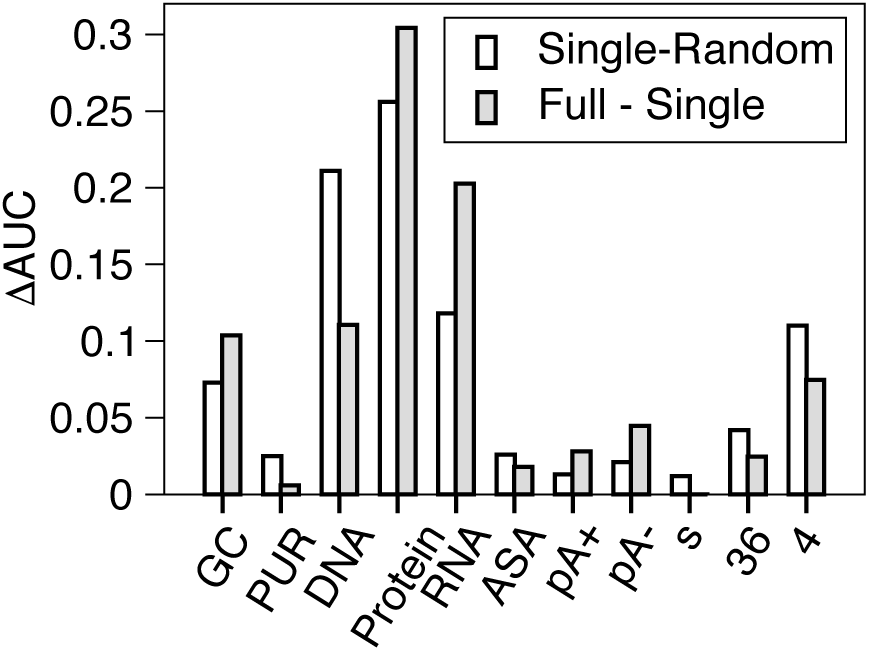
The difference in Area Under the ROC Curve (AUC) as a single feature group. Here, GC denotes GC content; PUR: Purine content; DNA: DNA conservation; Protein: Protein conservation; RNA: RNA conservation; ASA: Accessible surface area; pA+: polyA+ RNA-seq; pA-: polyA-RNA-seq; s: small RNA-seq; 36: H3K36me3 modification; and 4: H3K4me3 modification. The different is multiplied by 10 for removing single feature only (filled bar), to facilitate comparison to the results of using single feature (open bar).

The importance of protein conservation found in the above analysis is likely due to the presence of mRNAs in the negative samples. To further explore the features important for separating EVlncRNAs from HTlncRNAs only, we employed the training set (799 EVlncRNAs as the positive set and 799 HTlncRNAs without mRNAs as the negative set) and obtained the result of ten-fold cross validation with the full feature model. Then, we removed the redundant single feature that led to the largest increase of the AUC value in each round until the AUC can no longer increase after feature removal. The final model eliminated three features (protein conservation, predicted RNA ASA, and purine content) that are not useful for discrimination against HTlncRNAs. The most important remaining features according to the magnitude in AUC reduction after removal are RNA conservation (ΔAUC = 0.025), followed by DNA conservation (ΔAUC = 0.007) and the experimental feature of H3K36me3 modification (ΔAUC = 0.006). The same trend is observed based on reduction in precision with a fixed sensitivity

### Discrimination against mRNAs

Our method was designed to discriminate EVlncRNAs against both HTlncRNAs and mRNAs as all of our negative sets in training and test sets contain 1:1 ratio of mRNA : HTlncRNAs. To further examine the capability of discrimination against mRNAs, we built an additional test set by using the EVlncRNAs in our test set as the positive set, and newly randomly selected mRNAs as an additional negative set. EVlncRNA-pred achieves a high AUC of 0.959, a precision of 0.987, a specificity of 0.991, and the MCC value of 0.675. This higher performance in mRNA discrimination is consistent with our intuition that separating EVlncRNAs from mRNAs is easier than separating them from HTlncRNAs.

### Comparison with other methods

To the best of our knowledge, the method reported here is the first technique for discriminating EVlncRNAs from HTlncRNAs and mRNAs. Existing techniques for lncRNA prediction are dedicated to separate HTlncRNAs from mRNAs. We do not expect that they could be useful for identifying EVlncRNAs from HTlncRNAs and mRNAs. To confirm this, Figure 4 reported the applications of the logistic regression model CPAT [30], the random forest model COME [34], and support vector machines models CNCI [36] and PLEK [38] to the human test set. Indeed, CNCI, CPAT, and PLEK methods are close to random predictions at low false positive rates whereas COME is unable to make any positive prediction until false positive rates are greater than 0.2. Overall prediction of COME, CNCI, CPAT, and PLEK is better than random with AUCs ranging from 0.567, 0.672, 0.699, and 0.569, respectively. This is because mRNA belongs to the negative set whereas EVlncRNAs belongs to the positive set in training COME, CNCI, CPAT and PLEK. We would like to emphasize that the comparison made in Figure 4 is not to illustrate the improvement of our method over previous techniques but to highlight the difference in the prediction goals.

**Figure 4.**
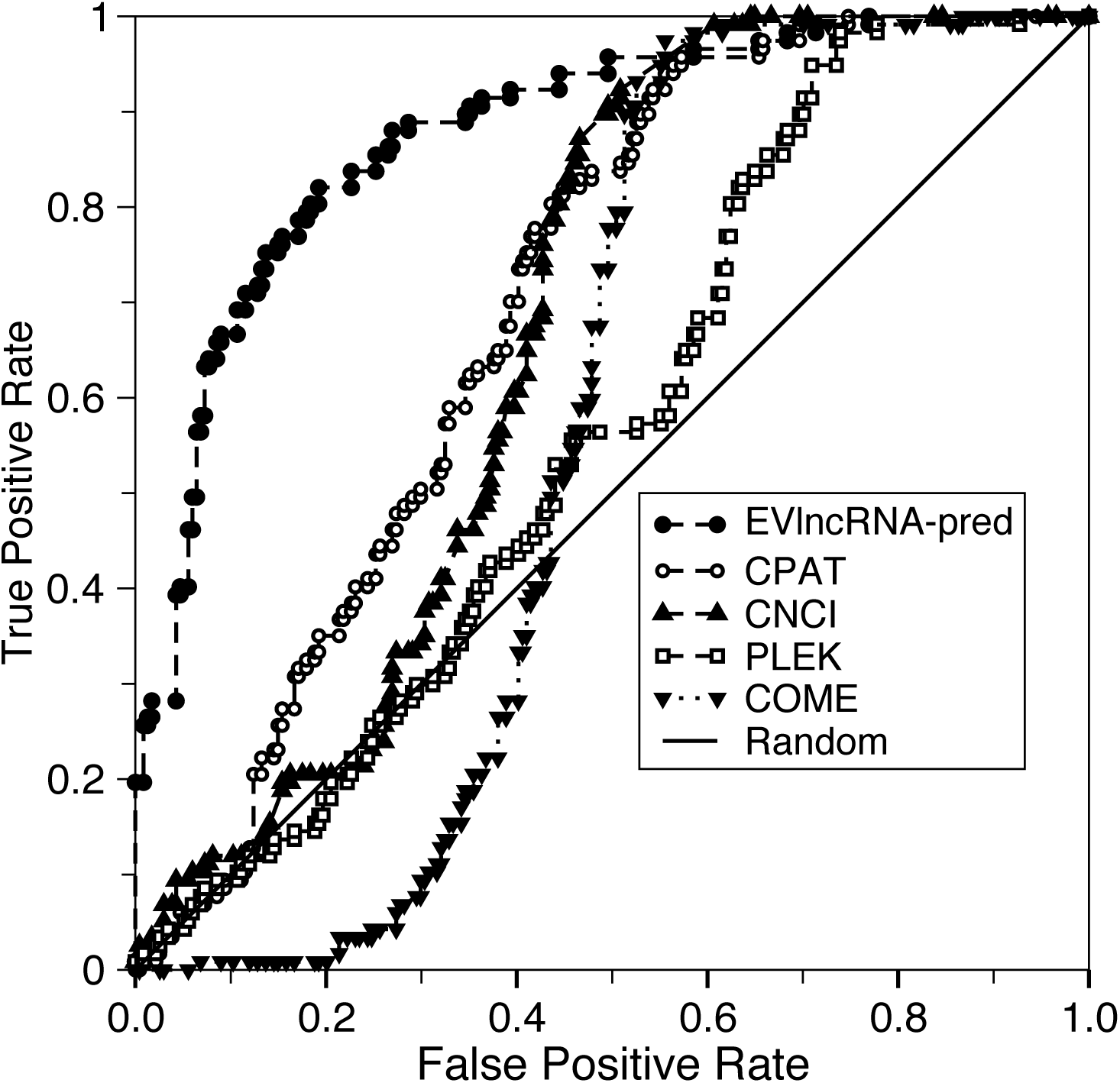
Receiver operating characteristic curves on the human test set by EVlncRNA-pred and several methods that were trained for separating expressed lncRNAs from mRNAs only.

### Case Studies

#### Tumour specific EVlncRNAs

Tumour-specific EVlncRNAs is a large group in all known EVlncRNAs. In the EVlncRNA database, there were 446 and 72 tumour-specific lncRNAs in our training and test sets, respectively. The sensitivity of EVlncRNA-pred (the fraction of predicted EVlncRNAs in known EVlncRNAs) is 54% for these tumour-specific lncRNAs in the training set and 57% in the test set, which are close to a sensitivity of 60% in ten-fold cross validation (Table 1), suggesting overall consistency of the method performance for specific types of EVlncRNAs.

#### CRISPRi-identified functional lncRNAs

Recently Liu *et al*. (2017) developed a CRISPR interference technique for a large-scale screening of lncRNA loci required for robust cell growth. Strictly speaking, the resulting 499 lncRNA loci discovered would require further validation by low-throughput experiments. However, it is of interest to examine the performance of EVlncRNA-pred for these newly discovered putatively functional lncRNAs. Among these 499 lncRNA loci, we located 194 lncRNAs with the gene structure information in the general transfer format, 59 lncRNAs of which are in the positive training set (known EVlncRNAs). Applying EVlncRNA-pred to the remaining 135 lncRNAs yields a sensitivity of 42%. It should be noted that the above putatively functional lncRNA loci were filtered with an experimental confidence score (called “screen score”) >7. If we increase this threshold from 7, 15 to 25, the sensitivity of our method will improve from 42%, 49%, to 67%. Concurrent increase of our method sensitivity and the experimental confidence score confirms the ability of EVlncRNAs to locate truly functional lncRNAs.

#### Newly discovered EVlncRNAs

We conducted a literature search for newly discovered EVlncRNAs because true positives from the EVlncRNA database are based on the literature prior to May 2016. We found that 24 new functional human lncRNAs are indeed classified as EVlncRNAs by EVlncRNA-pred. As shown in Supplementary Table S1, these new lncRNAs were experimentally validated by low-throughput techniques such as qRT-PCR, western blot, and knockdown. Their functions range from microRNA and protein binding to expression regulation although not all new lncRNAs have a clearly identified molecular-level function. For example, Li *et al.* [46] found that a lncRNA, SNHG20, has a significantly higher expression in Colorectal Cancer (CRC) tissues than in corresponding normal tissues from 107 CRC patients. SNHG20 regulated cell growth through modulation of a series of cell cycle-associated genes. Similarly, Lu *et al*. [47] found that a higher expression level of a lncRNA, SOX21-AS1, positively correlated with the tumour size and the advanced stage of tumor-node-metastasis (TNM), and the inhibition of SOX21-AS1 induced p57 expression. SNHG20 and SOX21-AS1 are classified as EVlncRNAs by EVlncRNA-pred.

## Discussion

We have developed a method termed EVlncRNA-pred for selecting potentially functional lncRNAs from expressing lncRNAs found in high-throughput sequencing. Two different versions of the method were developed: one requires sequence information only whereas the other needs high-throughput experimental data in expression and histone modification. The results show that both versions can provide reasonably accurate separation of EVlncRNAs from HTlncRNAs and mRNAs, whereas the experimental data can provide additional improvement from 0.47 to 0.51 for Matthews correlation coefficient in ten-fold cross validation. The method trained by human RNAs is robust as it performs equally well in mouse RNA discrimination and to a lesser extent in plant RNA discrimination.

In this work, we have randomly chosen 799 HTlncRNA and mRNA sequences to match in number to the largest training set we currently have for EVlncRNAs. The equal number was chosen to maximize the learning [48]. To confirm the randomness for the choice of 799 HTlncRNAs and mRNAs, we have randomly selected 9 additional sets of 799 HTlncRNAs and mRNAs. The results of 10 fold-cross validations for the 10 sets are 0.475±0.015 for MCC and 0.842±0.01 for AUC. These small standard deviations indicate unbiased choices of negative sets. Furthermore, to confirm the usefulness of setting the ratio to 1, we systematically expanded the training set by increasing the ratio from 1 to 1.5, 2, 3, and 4. We found that there is a reduction of AUC values from 0.879 to 0.866, 0.808, 0.790, and 0.799, respectively, for the human independent test set as the ratio increases. We also built another test set with ratio of 1:4:4 for EVlncRNA: HTlncRNA: mRNA. We observed similar reduction of AUC values for this larger test set (from 0.864 to 0.855, 0.799, 0.785, and 0.792, respectively). Thus, the model trained by the data with the ratio of 1 has the best performance not only for the test set with the ratio of 1 but also for the ratio of 1:4:4.

One revealing fact is that the most discriminating features are related to conservations at protein levels followed by RNA levels. It turns out that protein conservation is the most important for separating from mRNAs whereas RNA conservation is the most important for separating from HTlncRNAs. This result provides the additional confidence for the method developed. Although sequence conservation signal for lncRNA is in general weak [49], it remains one essential feature for functional lncRNAs [50-53]. The result reported here indicates that the conservation signal can be picked up by a machine learning technique to highlight the intrinsic difference between those experimentally validated lncRNAs and those somehow expressed in high-throughput sequencing.

Here, we assumed from the outset that all lncRNAs reported in GENCODE are negative samples after excluding experimentally validated ones. This assumption was made despite the training set may contain a significant number of false negatives, which are true functional lncRNAs yet to be validated by low throughput experiments. If the majority of the presumed negatives were false negatives, one would not be able to develop a method to separate positive from negatives during training. The fact that a highly robust method can be made indicates that false negatives are not dominant and there is a population in HTlncRNAs separable from known EVlncRNAs. The existence of such a population in HTlncRNAs distinct from known EVlncRNAs itself is interesting, as transcriptional noise could be a source for some of the lncRNAs found by high-throughput experiments [18].

Using a negative set containing some false negatives is a common practice in machine learning because negatives are always more difficult to prove. For example, in studying pathogenic genetic variations, genetic variants found in 1000 genome projects on healthy individuals [54] are considered as neutral (non-disease causing) [55]. However, this assumption may not be correct for some late-onset disorders, in particular. It was shown that removing potential false negatives (the genetic variants with low minor allele frequency and potentially pathogenetic [56]) reduces the performance of the method trained. This suggests that having more data is more important than reducing potential false negatives in the training set [55].

To further examine the effect of potential errors in negatives, we randomly added 5% or 10% errors to nine-folds in the training set by assigning HTlncRNAs to EVlncRNAs and EVlncRNAs to HTlncRNAs and testing the method for the remaining fold. This was repeated 10 times (ten-fold cross validation). We also randomly selected 5% or 10% errors 10 separate times to obtain an average effect. Introducing 5% and 10% errors leads to the average MCC values changed only slightly from 0.513 to 0.496 and 0.480, respectively. The small changes due to assignment errors indicate that our method is robust against potential assignment errors in the training set. However, one has to be cautious that not all positive predictions are functional lncRNAs as the fraction of correct predictions in positive predictions is at 81% for the human test set (i.e. 19% are incorrect). Moreover, the coverage of functional lncRNAs (sensitivity) is at 64% due to the small training set. That is, predictions may miss many functional RNAs, tissue-specific lncRNAs, in particular. Nevertheless, the method should be already useful for prioritizing potentially functional lncRNAs for further experimental validation. At the meantime, we hope to further improve sensitivity and precision in near future when a much larger dataset is available for deep learning.

To estimate the fraction of potentially functional EVlncRNAs in HTlncRNAs, we randomly selected 2000 human lncRNAs in NONCODE database [57]. Among them, 566 lncRNAs were classified as EVlncRNAs by EVlncRNA-pred. This would place the fraction at 28.3% (566/2000) in expressed lncRNAs found by high-throughput sequencing that possess the same characteristics as EVlncRNAs according to our machine-learning method. This indicates that many functional lncRNAs are waiting to be validated.

## Conclusions

In summary, known experimentally validated lncRNAs are separable from the majority of lncRNAs and mRNAs found in high-throughput experiments. The classification performance of the method EVlncRNA-pred is reasonably high with AUC >0.84 for independent tests on human and mouse RNAs and >0.8 for plant RNAs. This indicates that the model built here is already useful for prioritizing functional lncRNAs for validation by low throughput experiments. The method along with the training and test datasets is freely available for experimental biologists at http://biophy.dzu.edu.cn/lncrnapred/index.html.

## Materials and methods

### Training and test datasets for human lncRNAs

Most previous methods for discriminating lncRNAs from mRNAs were trained by using lncRNAs from GENCODE [43] as the positive dataset. These lncRNAs were obtained from the ENCODE project [58] by using a variety of high-throughput techniques and annotated by a combination of computational analysis, sequence comparison and manual annotation. Here we treated them as the negative dataset after excluding all known experimentally validated, functional lncRNAs from a recently curated database EVlncRNAs [16] (the positive dataset). Because EVlncRNAs is far from a complete dataset for functional lncRNAs, our negative dataset likely contains some false negatives. As we discussed in the discussion section, this should not prevent us addressing the question if current experimentally validated lncRNAs (denoted as EVlncRNAs for convenience) are separable from lncRNAs from high-throughput (HT) experiments (denoted as HTlncRNAs for convenience).

We first created a positive human test set from EVlncRNAs that were not contained in GENCODE V19 so that we can create a set of newly discovered, experimentally validated lncRNAs. This test set was obtained by using CD-HIT [59] to remove redundant sequences with more than 80% sequence similarity with HTlncRNAs in GENCODE V19 and among themselves. We have chosen 80% sequence identity cutoff because statistics suggests a significant reduction in secondary structure similarity for RNA sequences with <80% sequence identity [60]. Moreover, it is the lowest sequence identity cutoff allowed by the program CD-HIT [59]. This cutoff was also employed previously for establishing non-redundant RNA sequences [61, 62]. A total of 117 human EVlncRNAs were obtained as an independent positive test set. The remaining human lncRNAs from EVlncRNAs were used to generate the positive training set after removing redundant sequences by CD-HIT from the independent test set and among themselves. This leads to a training set of 799 human EVlncRNAs. The negative sets for HTlncRNAs (799 training and 117 independent test HTlncRNAs) were randomly selected from GENCODE V19 while ensuring <80% sequence identity among themselves and from the positive sets.

In addition to the HTlncRNA set as the negative set, we also included mRNAs from GENCODE V19 as the negative set. These mRNAs were randomly selected with <80% sequence similarity between each other and from selected HTlncRNAs and EVlncRNAs. The number of mRNAs is set to the same as the size of the two positive sets. Thus, the final human training dataset contains 799 EVlncRNAs (positive), 799 HTlncRNAs (negative) and 799 mRNAs (negative). The final human independent test set contains 177 EVlncRNAs (positives), 177 HTlncRNAs (negatives) and 177 mRNAs (negatives). Using both HTlncRNAs and mRNAs in the negative sets is to ensure that our method can discriminate EVlncRNAs from either HTlncRNAs or mRNAs.

Here, we have set the ratio of EVlncRNA:HTlncRNA:mRNA to 1:1:1 in training and test sets. The purpose is to maximize the learning by undersampling the negative samples [48]. To examine the effect of the ratio, we also built the training set for EVlncRNA:HTlncRNA:mRNA at 799:1200:1200 (1:1.5:1.5), 799:1600:1600 (1:2:2), 799:2400:2400 (1:3:3), and 799:3200:3200 (1:4:4), respectively. These additional mRNAs and HTlncRNAs were randomly selected with <80% sequence similarity between each other and from previously selected mRNAs, HTlncRNAs and EVlncRNAs. Similarly, we built an additional independent test set EVlncRNA:HTlncRNA:mRNA at the ratio of 117:468:468 (1:4:4) with the same positive samples in the independent test set but expanded the sample sizes of HTlncRNAs and mRNAs. A ratio of 1:4 between EVlncRNA and HTlncRNA is close to the real-world situation as we shall see.

### Independent test sets from mouse and plant lncRNAs

To further test the robustness of the methods developed, we established independent test sets by using mouse and plant lncRNAs. Similar to human datasets, the positive sets for plant and mouse were obtained from the EVLncRNAs database [16]. There were 166 mouse EVlncRNAs after removing redundant sequences with more than 80% sequence similarity to the human set (both positive and negative sets) and among themselves. We randomly selected 166 mouse HTlncRNAs from GENCODE V19 as the negative set after removing the redundant sequences from the mouse positive set, the human set and among themselves. In addition, we have randomly selected 166 mouse mRNA set from GENCODE V19 with the same sequence similarity cutoff to remove redundancy. The final mouse test set contains 166 EVlncRNAs (positives), 166 HTlncRNAs (negatives) and 166 mRNAs (negatives).

We used the EVlncRNAs of *Arabidopsis thaliana* in the EVLncRNAs database [16] to construct the positive set for plant. After removing redundant sequences with more than 80% sequence similarity with the human set and among themselves, 120 *Arabidopsis thaliana* EVlncRNAs were obtained as the plant positive set. The HTlncRNAs and mRNAs of *Arabidopsis thaliana* were from the Ensembl Plants database [63]. Similar to the mouse negative set, equal number of HTlncRNAs and mRNAs of *Arabidopsis thaliana* were randomly selected after removing the redundant sequences from the plant positive set, the human set, and among themselves. The final plant test set contains 120 EVlncRNAs (positives), 120 HTlncRNAs (negatives) and 120 mRNAs (negatives).

### Input features

All features were block-averaged similar to previous studies [34, 36, 38, 64]. Each block has 100 nucleotides, centered at 50, 100, 150, and etc. until the entire sequence is covered by blocks. For a given feature, the value or averaged value of each block was calculated to represent the block. The final feature values are average, maximum and variance of values of the blocks that covered the entire sequence. Following features are calculated.

#### Features based on sequences

Features based on sequences are employed to develop the sequence-only model.

#### GC content

The percentage of G and C in a sequence block was calculated.

#### Purine (PUR) content

The percentage of purines (G and A) in a sequence block was calculated.

#### DNA conservation score

The phastCons [44] scores provided by the UCSC [45] represented the DNA sequence conservation of human and mouse. The phastCons scores for human are from phastCons100way, and the scores for mouse are from phastCons60way. However, no similar scores are available for plant. We have simply set these values to zero when applying to the plant set (also see below).

#### Protein conservation score

The protein conservation score was calculated by BLASTx that searches a given nucleotide sequence against the protein sequence in the UniProt database [65].

#### RNA conservation score

Infernal (“INFERence of RNA Alignment”) [66] was employed for searching Rfam databases [67] for RNA structure and sequence similarities. It is an implementation of a special case of profile stochastic context-free grammars called covariance models (CMs). A CM is like a sequence profile, but it scores a combination of sequence conservation and RNA secondary-structure conservation.

#### Predicted solvent accessible surface area (ASA) of RNA

RNA ASA values were predicted by RNAsnap [61].

#### Features based on high-throughput experimental results

The above sequence-based features together with the features based on high-throughput experimental results (see below) are utilised to develop the full-feature model.

#### Expression abundance

The reads per kilobase per million (RPKM) for each sequence were calculated from polyA+, polyA- and small RNA-seq data. The maximum scores for five cell lines (GM12878, K562, H1-hESC, HeLa-S3, and HepG2) were assigned to human sequence. For the mouse and *Arabidopsis thaliana*, the RNA-seq data of various tissues were used. These feature values were obtained from COME [34].

#### Histone modification

The ChIP-seq data from H3K36me3 and H3K4me3 modification were used to calculate the signal over a sequence. The averaged input-normalized signals of five cell lines (GM12878, K562, H1-hESC, HeLa-S3, and HepG2) were used for human sequences. The ChIP-seq data of various tissues were used for sequences of mouse and *Arabidopsis thaliana*. These values were obtained from COME [34].

Ribosome profiling is another possible experimental feature. However, a previous study suggested its minor contribution to discrimination of HTlncRNA from mRNA [34]. We expect that it is less useful for separating EVlncRNA from HTlncRNA as both are not translated into peptides or proteins. As a result, this feature was not employed in this study.

### Support vector machines

We used SVM with the RBF kernel implemented in LIBSVM version 3.22 [68] to build our model. We optimized the parameters C and gamma using the grid search algorithm implemented in LIBSVM.

### Cross validation and independent test

We performed 10-fold cross validation on the training set. In this cross validation, the training set was randomly divided into ten folds, and each fold was tested in turn by using the remaining nine folds for training. To examine whether the results are consistent for different divisions of the dataset, we conducted 10-fold cross validations 100 times by randomly dividing the training set 100 times. We also used the whole training set to train the model and tested the model on independent test sets.

### Performance evaluation criteria

The performance of our method was evaluated by Matthews correlation coefficient (MCC), receiver operating characteristic (ROC) curve, area under the ROC curve (AUC), accuracy, sensitivity, specificity, and precision. The equations are as below.

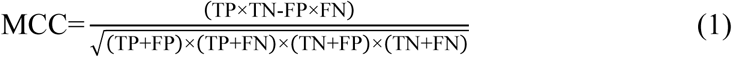

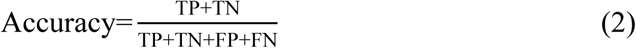

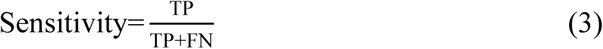

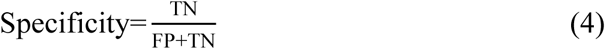

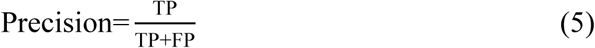

where TP and TN represent the positive and negative samples that have been correctly predicted, respectively, FP and FN represent the positive and negative samples that have been falsely predicted, respectively. MCC is essentially a correlation coefficient between predicted and actual binary classifications with values between −1 to 1 with zero for random prediction. It is a balanced measure for unequal-sized positive and negative samples. Sensitivity is the fraction of predicted true EVlncRNAs in all true EVlncRNAs. Specificity is the fraction of predicted true negatives in all true negatives. Precision is the fraction of true EVlncRNAs in all predicted EVlncRNAs.

### Data and software availability

EVlncRNA-pred is available as a web server at http://biophy.dzu.edu.cn/lncrnapred/index.html. All datasets used in this study can be obtained from the same website.

## Acknowledgements

We thank Hucheng Tang for helping develop the web-based server.

## Disclosure statement

No potential conflict of interest was reported by the authors.

## Funding

This work was supported by the National Natural Science Foundation of China [61671107, 61271378, 61801081]; Taishan Scholars Program of Shandong province of China [Tshw201502045]; National Health and Medical Research Council of Australia [1121629 to Y.Z.]; Australia Research Council [DP 180102060 to Y.Z.]; and Talent Introduction Project of Dezhou University of China [320111 to B.Z.].

