## Supplemental Table S1 for "Predicting functional long non-coding RNAs validated by low throughput experiments"

*[a]Shandong Provincial Key Laboratory of Biophysics, Institute of Biophysics, Dezhou University, Dezhou, China; [b]College of Physics and Electronic Information, Dezhou University, Dezhou, China; [c]School of Data and Computer Science, Sun Yat-sen University, Guangzhou, China; [d]Institute for Glycomics and School of Information and Communication Technology, Griffith University, Gold Coast, QLD, Australia*

*corresponding author: Yaoqi Zhou,, address: Institute for Glycomics and School of Information and Communication Technology, Griffith University, Gold Coast, QLD 4222, Australia; Jihua Wang,, address: Shandong Provincial Key Laboratory of Biophysics, Institute of Biophysics, Dezhou University, Dezhou 253023, China

### SUPPLEMENTAL TABLE

Table S1. Newly validated human lncRNAs.

| Name | Function | Method | Ref. |
|---|---|---|---|
| ZNF582-AS1 | Suppress colony formation by CRC cells | qRT-PCR, ectopic expression | [1] |
| SOCS2-AS1 | Modulating the epigenetic control | qRT-PCR, knockdown, Western Blot, ChIP Assay, RIP Assay | [2] |
| SNHG20 | Promote cell proliferation, etc. | qRT-PCR, western blot, knockdown | [3] |
| SNHG18 | Inhibit semaphorin 5A expression | qRT-PCR, knockdown, in situ hybridization, immunohistochemistry, Western blot | [4] |
| LNCPRESS1 | Interacts with SIRT6 | qRT-PCR, ChIP-qPCR | [5] |
| MHENCR | Bound to miR-425 and miR-489 | qRT-PCR, knockdown, Western blot, RNA Immunoprecipitation | [6] |
| SATB2-AS1 | Expression regulation | qRT-PCR, knockdown | [7] |
| PTOV1-AS1 | Expression regulation | knockdown, western blotting and Ago2 immunoprecipitation | [8] |
| LINC00441 | Epigenetic interaction with RB1 | gain- and loss-of-function investigation, RNA pull-down assay, Luciferase reporter gene assay, Chromatin immunoprecipitation | [9] |

| | | | |
|---|---|---|---|
| LINC01013 | Induce snail | knockdown, overexpression | [10] |
| HCG11 | Interaction with IGF2BP1 | knockdown, Western blot | [11] |
| TPM1-AS | Regulates the alternative splicing of TPM1 | In situ hybridization and RNA immunoprecipitation assays, knockdown, overexpression, qRT-PCR, western blot | [12] |
| LINC00319 | Binding with miR-32 | knockdown, Luciferase acitivity and RNA pull-down assays | [13] |
| SOX21-AS1 | Expression inhibition | qRT-PCR, western-blot and immunohistochemistry | [14] |
| ZNF503-AS1 | Downregulation of ZNF503 expression. | qRT-PCR, Fluorescent in situ hybridization, Immunoblotting, | [15] |
| LINC00968 | Oncogene in non-small cell lung cancer | qRT-PCR, Western Blot, knockdown, overexpression | [16] |
| SNHG17 | Binding to enhancer of zeste homolog 2 | qRT-PCR, knockdown, RNA immunoprecipitation (RIP) assay, Chromatin immunoprecipitation (ChIP) assay, Western blot | [17] |
| FIRRE | Interacts with heterogeneous nuclear ribonucleoproteins U | qRT-PCR, knockdown, Western blot, Northern blot, RNA immunoprecipitation, luciferase assay | [18] |

| | | | |
|---|---|---|---|
| GHRLOS | Cancer biomarker | qRT-PCR | [19] |
| MIR22HG | Cancer prognosis | qRT-PCR, knockdown | [20] |
| IDH1-AS1 | Repressed by c-Myc | knockdown, overexpression | [21] |
| FOXD3-AS1 | Interaction with miR-150 | Northern blot, knockdown, overexpression, qRT-PCR, Western blot, Luciferase reporter assay | [22] |
| LINP1 | Facilitates DNA damage repair through non-homologous end joining (NHEJ) pathway | qRT–PCR, knockdown, RNA immunoprecipitation, western blot, Fluorescence in situ hybridization | [23] |
| MIR222HG | Association with the RNA-induced silencing complex | qRT-PCR, Dual-luciferase reporter assay, knockdown, overexpression | [24] |

[7] Liu SH, Zhu JW, Xu HH, et al. A novel antisense long non-coding RNA SATB2-AS1

overexpresses in osteosarcoma and increases cell proliferation and growth. Mol Cell Biochem. 2017;430:47-56.

[13] Zhou B, Yuan W, Li X. Long intergenic noncoding RNA 319 (linc00319) promotes cell proliferation and invasion in lung cancer cells by directly downregulating the tumor suppressor MiR-32. Oncol Res. 2017.

[23] Wang X, Liu H, Shi L, et al. LINP1 facilitates DNA damage repair through non-

homologous end joining (NHEJ) pathway and subsequently decreases the sensitivity of cervical cancer cells to ionizing radiation. Cell cycle. 2018;17:439-447.

[24] Sun T, Du SY, Armenia J, et al. Expression of lncRNA MIR222HG co-transcribed from the miR-221/222 gene promoter facilitates the development of castration-resistant prostate cancer. Oncogenesis. 2018;7:30.
